# Effects of vestibular stimulation on gait stability when walking at different step widths

**DOI:** 10.1101/2021.09.09.459650

**Authors:** Rina M. Magnani, Jaap H. van Dieën, Sjoerd M. Bruijn

**Author notes:** Corresponding author: Rina Marcia Magnani, Rua T-53, 1770, 1206, 74150-310. Goiânia/GO, Brazil.

## Abstract

Vestibular information modulates muscle activity during gait, presumably to contribute to stability. If this is the case, stronger effects of perturbing vestibular information on local dynamic stability of gait, a measure of the locomotor system’s response to small, naturally occurring perturbations, can be expected for narrow-base walking (which needs more control) than for normal walking and smaller effects for wide-base walking (which needs less control). An important mechanism to stabilize gait is to coordinate foot placement to center of mass (CoM) state. Vestibular information most likely contributes to sensing this CoM state. We, therefore, expected that stochastic electrical vestibular stimulation (EVS) would decrease the correlation between foot placement and CoM state during the preceding swing phase. In fourteen healthy participants, we measured the kinematics of the trunk (as a proxy of the CoM), and feet, while they walked on a treadmill in six conditions: control (usual step width), narrow-base, and wide-base, each with and without stochastic EVS (peak amplitude of 5 mA; RMS of ^~^ 1.2 mA; frequency band from 0 to 25 Hz). Stochastic EVS decreased local dynamic stability irrespective of step width. Foot placement correlated stronger with trunk motion during walking with EVS than without in the control condition. However, residual variance in foot placement was increased when walking with EVS, indicating less precise foot placement. Thus, a vestibular error signal leads to a decrease in gait stability and precision of foot placement but these effects are not consistently modulated by step width.

## Introduction

Gait stability has been defined as the ability to walk without falling, in spite of intrinsic or environmental perturbations (Bruijn et al. 2013). Gait stability is most challenged in the mediolateral direction (Bauby and Kuo 2000), due to the narrow base of support in this direction, and the movement towards the lateral edge of the base of support during each single-leg stance phase. It is assumed that mediolateral gait stability requires feedback control based on sensory information (Bauby and Kuo 2000).

Electrical vestibular stimulation (EVS) has been applied to probe the role of vestibular feedback in the control of standing and walking. The stimulus causes artificial stimulation of the afferents of the semicircular canals and otoliths, resulting in a superposition of error signals on top of a natural motion signal encoded by the vestibular system, leading to changes in perceived orientation and motor reactions (St George and Fitzpatrick 2011). In studies in standing humans, Galvanic binaural and bipolar stimuli at amplitudes from 0.5 to 1.2mA cause a shift of the center of pressure (CoP) towards the anodal side (Day et al. 1997; Yang et al. 2015; Matsugi et al. 2020), while during overground walking, similar stimuli lead to deviations in heading towards the anodal side (Fitzpatrick et al. 1999; Jahn et al. 2000; Bent et al. 2004). Stochastic, zero-mean rapidly fluctuating EVS is not expected to cause a prolonged illusion of motion in one direction, but it would increase the error in the afferent vestibular signal (Dakin et al. 2010), which likely perturbs feedback control of gait stability.

The local divergence exponent is an often used measure of gait stability (Bruijn et al. 2013). It represents the rate of divergence between neighboring trajectories in state space over a short time interval and indicates resistance to small perturbations (Dingwell and Cusumano 2000; Van Emmerik et al. 2016). The local divergence exponent has been shown to be associated with fall risk in several studies (Toebes et al. 2012; Rispens et al. 2014; van Schooten et al. 2016). Stochastic EVS has been shown to decrease gait stability (i.e. increase the local divergence exponent) (Sloot et al. 2011; van Schooten et al. 2011; Magnani et al. 2021).

Recently, the step-by-step relationship between foot placement and the center of mass (CoM) state has been proposed as a reflection of how gait is stabilized (Wang and Srinivasan 2014; Bruijn and Van Dieën 2018). The mediolateral CoM state during swing, or closely related pelvis or trunk states, predicts > 80% of the variance in mediolateral foot placement at the end of swing. For instance, when the CoM is positioned more medial relative to the stance leg, or when it is moving faster medially than on average during the swing phase, a more lateral foot placement than average at the end of this swing phase results, and vice versa. These adjustments in foot placement provide negative feedback, as a more lateral foot placement induces a larger medial acceleration of the center of mass and vice versa. The strength of the relation between CoM state and foot placement can thus be seen as an index of the quality of CoM feedback control. This quality would be dependent on the accuracy and precision of sensory information on the CoM state, which is likely based on multiple sensory modalities including vision, proprioception, and vestibular information (Bruijn and Van Dieën 2018). At gait initiation, vestibular stimulation specifically was shown to affect foot placement, suggesting that it indeed affects the estimate of the CoM movement used for foot placement control (Reimann et al. 2017). Thus, we expected that walking with stochastic EVS, would decrease the degree of foot placement control and consequently reduce local dynamic stability of walking.

Step width adjustments are commonly needed in daily life, to deal with environmental constraints, such as avoiding a puddle or a hole. Moreover, a wider step width has been reported as a response to impaired lateral stability during locomotion (Kubinski et al. 2015; Aboutorabi et al. 2016). However, previous studies have also shown that local dynamic stability is reduced when walking with wider steps (Young and Dingwell 2012; Magnani et al. 2021). Walking with wide steps requires less tight control (Perry and Srinivasan 2017), allowing people to be less stable (i.e. have a higher local divergence exponent), and stability may thus be less affected by disturbances of the vestibular information, as this information may simply be used less. At the same time, narrow-base walking, due to the challenge it poses to gait stability, is associated with more tightly controlled CoM motion and foot placement (Arvin et al. 2016; Perry and Srinivasan 2017) and this may suggest increased reliance on sensory, including vestibular, information. In line with this, vestibular evoked responses in ground reaction force and muscles activity are increased in magnitude when one is exposed to a postural threat (Horslen et al. 2014; Naranjo et al. 2015). In a recent study (Magnani et al. 2021)., we showed that, compared to normal walking, mediolateral ground reaction forces were less coupled to vestibular stimulation when subjects were stabilized by means of elastic cords providing conservative forces to maintain the mediolateral position of the body CoM or when walking with wide steps. Moreover, when walking with narrower steps, a stronger coupling between ground reaction force and EVS was present. Overall, these findings indicate that the importance of vestibular input for control of human walking is dependent on stability demands.

In the current study, we tested how stochastic EVS affects gait stability during normal walking, as well as during walking with imposed narrow and wide steps. We assessed the effect of EVS on gait stability by means of the local divergence exponent, as done in previous studies (Sloot et al. 2011; van Schooten et al. 2011) and on foot placement, suggested to be the main mechanism to regulate mediolateral stability (Bruijn and Van Dieën 2018). We hypothesized that stochastic EVS would increase the local divergence exponent and variability of trunk movement, decrease the amount of variance in foot placement that can be explained by CoM state (i.e. decrease R^2^), and increase the residual variance in foot placement. Moreover, we hypothesized that, given the dependence of the role of vestibular information on stabilization demands, these effects would be stronger in narrow-base walking than in control walking and less strong in wide-base walking.

## Methods

### Ethics Statement and Participants

We measured 23 healthy young adults between 24 and 33 years recruited from the university campus. Nine participants were excluded during data analysis due to technical problems in the collection of kinematic data in any of the trials recorded, as a consequence of construction work in the lab building. Data of fourteen healthy participants (five female/nine male; mean age 28.42±2.87 years) were included in the analysis. None of the participants reported any auditory, vestibular, neurologic, cardiovascular, or orthopedic disorders, and all had a normal or corrected-to-normal vision. The participants agreed to participate in the study by signing the informed consent form, and the study was approved by the Ethics Committee of the Faculty of Behavioural and Movement Sciences of the VU Amsterdam (VCWE-2017-158).

### Instrumentation

Kinematic data were recorded using a 3D motion analysis system (Optotrak Northern Digital Inc., Waterloo, Ontario, Canada) operating at 100 samples per second. Light emitting diodes (LED) clusters were positioned at the occipital lobe, the spinous process of the sixth thoracic vertebra (T6), the posterior superior iliac spines and bilaterally at the calcaneus. Forces were collected from force plates embedded in the treadmill (Motekforce Link, The Netherlands) sampling at 200 samples per second. Continuous percutaneous bipolar electrical stimulation was used to modulate the firing rate of the vestibular nerves, using an isolated linear stimulator (STMISOLA, BIOPAC System Inc., USA). When the head is facing forward, galvanic binaural stimulation evokes an illusion of head roll rotational velocity (Peters et al. 2015) about an axis directed posteriorly and superiorly by 18 degrees relative to Reid’s plane (Fitzpatrick and Day 2004; St George and Fitzpatrick 2011), and results in a postural response in the frontal plane to compensate for the induced roll error signal (Forbes et al. 2016; Tisserand et al. 2018). Here, we used bandwidth-limited stochastic EVS with a frequency content of 0 to 25 Hz. This stimulus bandwidth and amplitude generally produces very little interpretable perceptual signal of motion. However, an analysis in which we band-pass filtered T6 marker position at the stride frequency showed that the standard deviation of the remaining signal was increased during the EVS trials, which suggests that our stimulation had some effect on perception of motion and did also increase motion in response to this. This stimulus was created using MATLAB software (MathWorks, Natick, MA, USA) (Figure 1) from zero-mean low-pass filtered (25 Hz cutoff, zero lag, fourth-order Butterworth) white noise, and had a peak amplitude of 5 mA and root mean square (RMS) of ~ 1.2 mA. All volunteers were exposed to the same stimulus during eight minutes while walking. The stimulation was applied by flexible carbon surface electrodes (about 9 cm^2^). The electrodes were coated with Spectra 360 electrode gel (Parker Laboratories, Fairfield, NJ, USA) and affixed with adhesive tape to the clean and dry skin of the participants’ mastoid processes, and further stabilized by an elastic headband.

**Figure 1:**
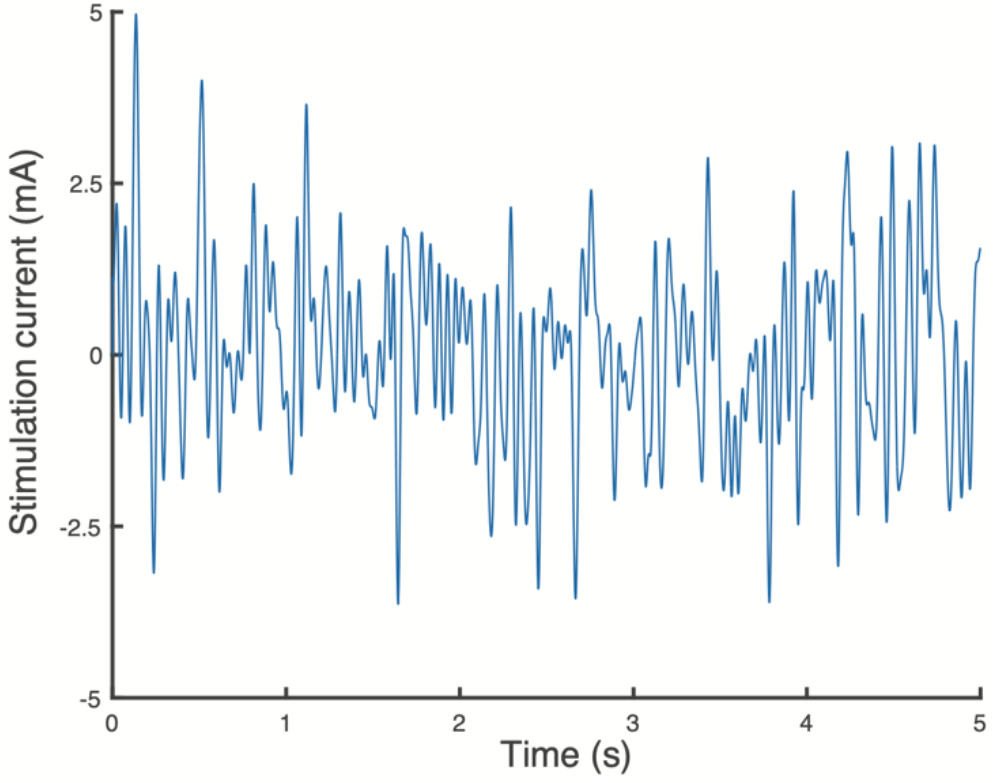
Vestibular stimulation signal.

### Protocol

Participants walked on a level treadmill (walking surface width of 102cm and length of 157cm) in six different conditions while wearing their own shoes. All conditions were performed at a fixed speed of 0.8 m/s, and cadence was imposed by the beat of a metronome, which was set at 78 steps/min. We controlled walking speed and cadence since both of these have been shown to influence vestibular contributions to locomotion, in such way that vestibular contributions to locomotion are highest at slow walking speeds and cadence (Dakin et al. 2013; Forbes et al. 2017). A walking speed of 0.8 m/s and cadence 78 steps/min were chosen to replicate the conditions used by Dakin et al. (2013). The conditions were characterized by step width, here called Condition (control, narrow-base, wide-base) and by the stimulus presence (EVS and no-EVS). For the control condition, participants walked with their usual step width; for narrow-base walking, participants were asked to adopt a smaller base of support (smaller than hip width); for wide-base walking participants were asked to increase the base of support adopting step width greater than hip width. The duration of each one of the EVS conditions (control, narrow and wide-base walking) was eight minutes, and the stimulus started with the measurement and was applied over all eight minutes of the trial. The no-EVS conditions took one minute. The order of the conditions was randomized for each participant, and participants were given a 2-minute break between conditions (with no stimulation or walking).

### Data analysis

Gait events were identified from center of pressure data (Roerdink et al. 2008), and a complete gait cycle was defined by the time between heel contact and subsequent heel contact of the same limb. Since the minimum number of strides over conditions and participants was 32, we calculated all gait parameters for 32 strides of each condition. We chose to analyze the last 32 strides, because in case of any habituation or reweighting as a consequence of EVS, we expected this to be the most stable part. We also performed the same analysis on the first 32 strides of each condition with similar results (results not reported here). The mean and standard deviation of step time (ST) (calculated as the duration between two consecutive heel contacts and step width (SW) (determined as the mediolateral distance between the heels during heelstrikes) were calculated. Gait stability was evaluated by the local divergence exponent calculated from the T6 mediolateral velocity time series (Dingwell and Cusumano 2000). To this end, the time series was first normalized to 3200 samples, after which a 5-dimensional state space was constructed from this time normalized signal and 4 time-delayed copies. We used a fixed delay of 10 samples, such that each dimension was shifted by 10 samples with respect to the previous one. We used a fixed number of dimensions and a fixed delay, as previous research has suggested that this yields the most sensitive and reliable results (van Schooten et al. 2013). Next, for each point in the state space, the divergence between the point and its nearest neighbor (defined as the point with the smallest Euclidean distance, while having at least half a cycle temporal separation to the original point), was tracked over time. The mean of the log of these curves was then taken as the logarithmic rate of divergence and the local divergence exponent was calculated as the slope to this curve from 0-0.5 strides (Rosenstein et al. 1993; Stenum et al. 2014; Bruijn 2017). In addition, we calculated CoM position variability at heelstrike, since trunk movement variability has previously been shown to increase due to EVS (van Schooten et al. 2011).

Foot placement control was quantified using a regression equation relating mediolateral foot placement to the position and velocity of mediolateral trunk CoM at heelstrike (Wang and Srinivasan 2014; van Leeuwen et al. 2020). The T6 marker was used to approximate the mediolateral CoM position. The mediolateral trunk velocity (VCoM) was calculated as the first derivative of the mediolateral trunk position time-series. We used the following regression equation:

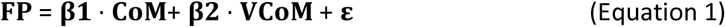

where β1 and β2 are the regression coefficients, *ε* the error (Wang and Srinivasan 2014; Mahaki et al. 2019; Bruijn and Van Leeuwen 2020; van Leeuwen et al. 2020), with all variables detected at heelstrike to focus on the outcome of the feedback process, which likely involves information obtained during the complete swing phase of gait. The ratio of predicted foot placement variance to actual foot placement variance at heelstrike (R^2^) was used as outcome measure. A higher R^2^ represents a stronger coupling between mediolateral trunk state and the subsequent mediolateral foot placement (Wang and Srinivasan 2014). However, as this R^2^ is a ratio of the variance in foot placement explained by the model and the total variance in foot placement, R^2^ will reflect coupling between trunk state and foot placement but also the magnitude of the variance in CoM state. This is especially problematic, as stochastic EVS has been shown to increase trunk movement variability. Thus, we also calculated the residual variance in foot placement with respect to the predicted foot placement (i.e. mean(| ε|).

### Statistical Analysis

The assumption of normality was checked by the Shapiro-Wilk test (p> 0.05). The effects of EVS and step width on SW, ST and their standard deviations, local divergence exponent (LDE), R^2^, residual variance of foot placement, and trunk position variability at heelstrike were tested using repeated measures analysis of variance with two factors, Stimulation (EVS – electrical vestibular stimulation and no-EVS) x Condition (control, narrow, wide). A Bonferroni correction was applied for the post hoc tests, which were only performed for narrow vs. control and wide vs. control. Whenever a significant interaction effect was found, we tested whether the change due to EVS was significant between conditions (i.e. we tested the paired difference of EVS-no EVS between conditions). All statistical analyses were performed in Matlab, using p< 0.05 as a threshold for significance.

## Results

The results of all repeated measures ANOVAs are presented in Table 1 and the results of the post hoc tests are presented in Tables 2 (effects of Condition) and 3 (interaction effects of Condition and EVS). In accordance with the instructions, Condition significantly affected step width, where step width was significantly wider during wide-base than control walking and significantly narrower in narrow-base compared to the control walking (Figure 2A). Step width showed no significant EVS effect or Condition x EVS interaction. Step width variability showed significant EVS and Condition effects, as well as a significant Condition x EVS interaction (Figure 2B). These effects indicated that step width variability increased when walking with EVS. Moreover, step width variability was smaller in the control condition than during narrow-base walking (Table 2). The interaction indicated a smaller effect of EVS for wide-base walking compared to the other conditions, but post-hoc tests (Table 3) were not significant.

**Table 1:** ANOVA results. Bold values represent statistically significant results (p<0.05). SW: step width; ST: step time; var: variability; LDE: local divergence exponent; R^2^: percentage of explained variance of foot placement; EVS: electrical vestibular stimulation.

|  | EVS | Condition | EVS x Condition |
| --- | --- | --- | --- |
| SW (m) | $F(1,13)=1.74$; $p=0.21$; $\eta_p^2=0.12$ | **$F(2,26)=49.28$; $p=0.00$; $\eta_p^2=0.79$** | $F(2,26)=2.68$; $p=0.09$; $\eta_p^2=0.17$ |
| SW var (m) | **$F(1,13)=15.27$; $p=0.00$; $\eta_p^2=0.54$** | **$F(2,26)=7.21$; $p=0.00$; $\eta_p^2=0.36$** | **$F(2,26)=6.00$; $p=0.01$; $\eta_p^2=0.32$** |
| ST (s) | **$F(1,13)=6.12$; $p=0.03$; $\eta_p^2=0.32$** | **$F(2,26)=4.51$; $p=0.02$; $\eta_p^2=0.26$** | **$F(2,26)=5.06$; $p=0.01$; $\eta_p^2=0.28$** |
| ST var (s) | **F(1,13)=11.24; p=0.01; $\eta_p^2=0.46$** | F(2,26)=2.17; p=0.13; $\eta_p^2=0.14$ | F(2,26)=0.11; p=0.90; $\eta_p^2=0.01$ |
| LDE | **F(1,13)=11.34; p=0.01; $\eta_p^2=0.47$** | **F(2,26)=19.99; p=0.00; $\eta_p^2=0.61$** | F(2,26)=0.02; p=0.98; $\eta_p^2=0.00$ |
| CoM var | **F(1,13)=33.17; p=0.00; $\eta_p^2=0.72$** | **F(2,26)=9.16; p=0.00; $\eta_p^2=0.41$** | F(2,26)=2.86; p=0.08; $\eta_p^2=0.18$ |
| R<sup>2</sup> | F(1,13)=2.61; p=0.13; $\eta_p^2=0.17$ | **F(2,26)=12.77; p=0.00; $\eta_p^2=0.50$** | **F(2,26)=5.57; p=0.01; $\eta_p^2=0.30$** |
| Residual variance | **F(1,13)=25.76; p=0.00; $\eta_p^2=0.66$** | **F(2,26)=5.02; p=0.01; $\eta_p^2=0.28$** | **F(2,26)=5.86; p=0.01; $\eta_p^2=0.31$** |

**Table 2:** P-values of the post-hoc comparisons of the walking Conditions. Bold values represent statistically significant results (p<0.05). SW: step width; ST: step time; var: variability; LDE: local divergence exponent; R^2^: percentage of explained variance of foot placement; EVS: electrical vestibular stimulation. Note: whenever a main effect of Condition in the ANOVA was not significant, corresponding post-hoc tests were not performed, hence the empty cells.

|  | Control-Narrow | Control-Wide | Wide-Narrow |
| --- | --- | --- | --- |
| SW (m) | 0 | 0 | 0 |
| SW var (m) | 0 | 0,46 | 0,296 |
| ST (s) | 0,089 | 1 | 0,099 |
| ST var (s) |  |  |  |
| LDE | **0,006** | 0,051 | 0 |
| CoM var | **0,001** | 1 | 0,05 |
| R<sup>2</sup> | **0,001** | 1 | **0,003** |
| Residual variance | **0,045** | 0,064 | 1 |

**Table 3:** P-values of the post-hoc comparisons of the effects of EVS between walking Conditions. Bold values represent statistically significant results (p<0.05). SW: step width; ST: step time; var: variability; LDE: local divergence exponent; R^2^: percentage of explained variance of foot placement; EVS: electrical vestibular stimulation. Note: whenever the interaction effect in the ANOVA was not significant, corresponding post-hoc tests were not performed, hence the empty cells.

| | $\Delta\text{EVS}(\text{Control}) - \Delta\text{EVS}(\text{Narrow})$ | $\Delta\text{EVS}(\text{Control}) - \Delta\text{EVS}(\text{Wide})$ | $\Delta\text{EVS}(\text{Wide}) - \Delta\text{EVS}(\text{Narrow})$ |
| --- | --- | --- | --- |
| SW (m) |  |  |  |
| SW var (m) | 1 | 0,06 | 0,054 |
| ST (s) | 0,323 | 0,266 | 0,052 |
| ST var (s) |  |  |  |
| LDE |  |  |  |
| CoM var |  |  |  |
| **R<sup>2</sup>** | 0,157 | **0,015** | 0,747 |
| **Residual variance** | 0,225 | 0,222 | **0,041** |

**Figure 2:**
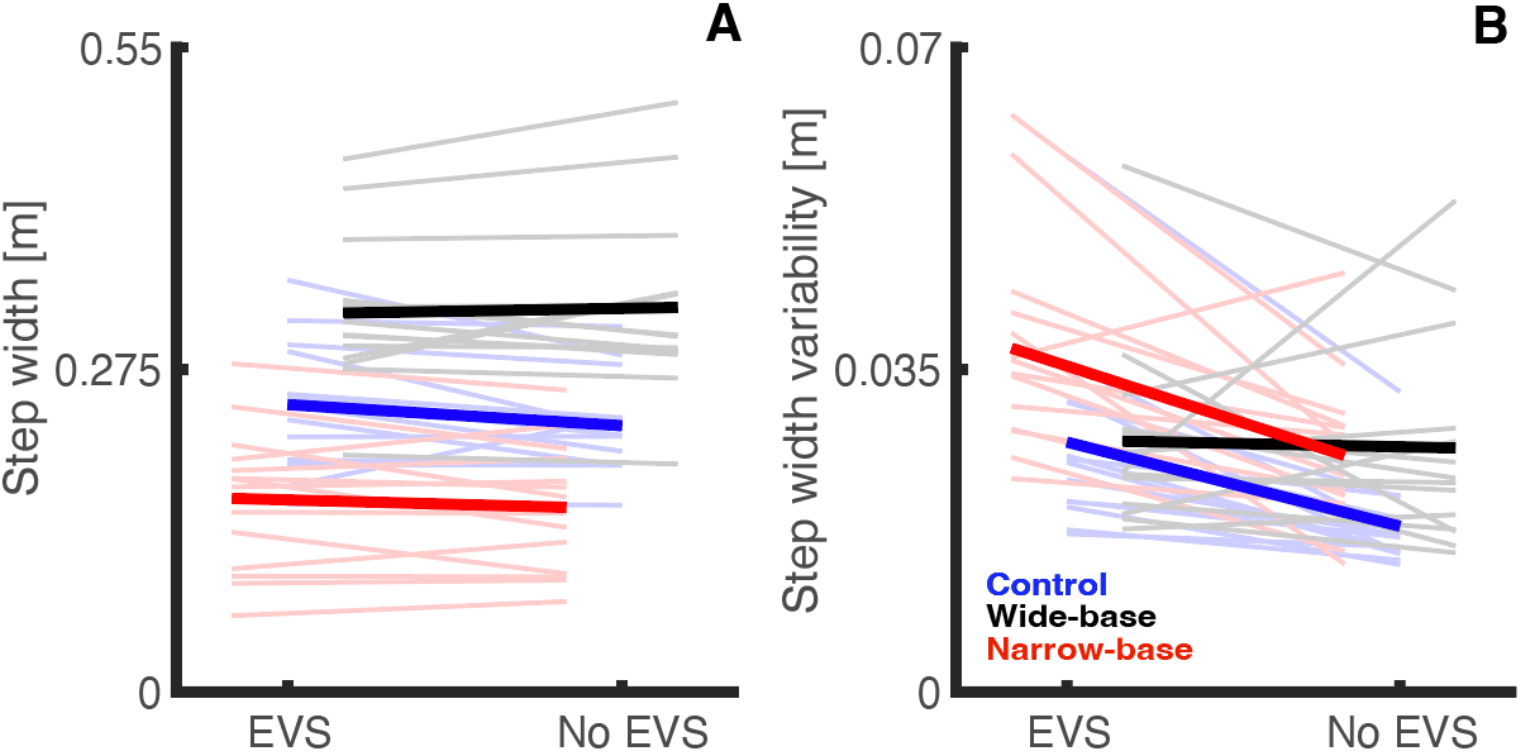
Effects of Condition and EVS on step width (A) and step width variability (B). Thick lines represent mean values and thin lines show individual data for each condition according to the following colors - blue lines: control condition; black lines: wide-base condition; red lines: narrow-base condition. Note: the conditions are jittered on the x-axis.

Step time was significantly higher during no EVS conditions (Figure 3A). In addition, step time showed a significant Condition effect, as well as a significant EVS X Condition interaction, but none of the post hoc test was significant. Step time variability was significantly higher during EVS conditions than during no-EVS conditions (Figure 3B) but was not affected by Condition or an interaction of EVS and Condition.

**Figure 3:**
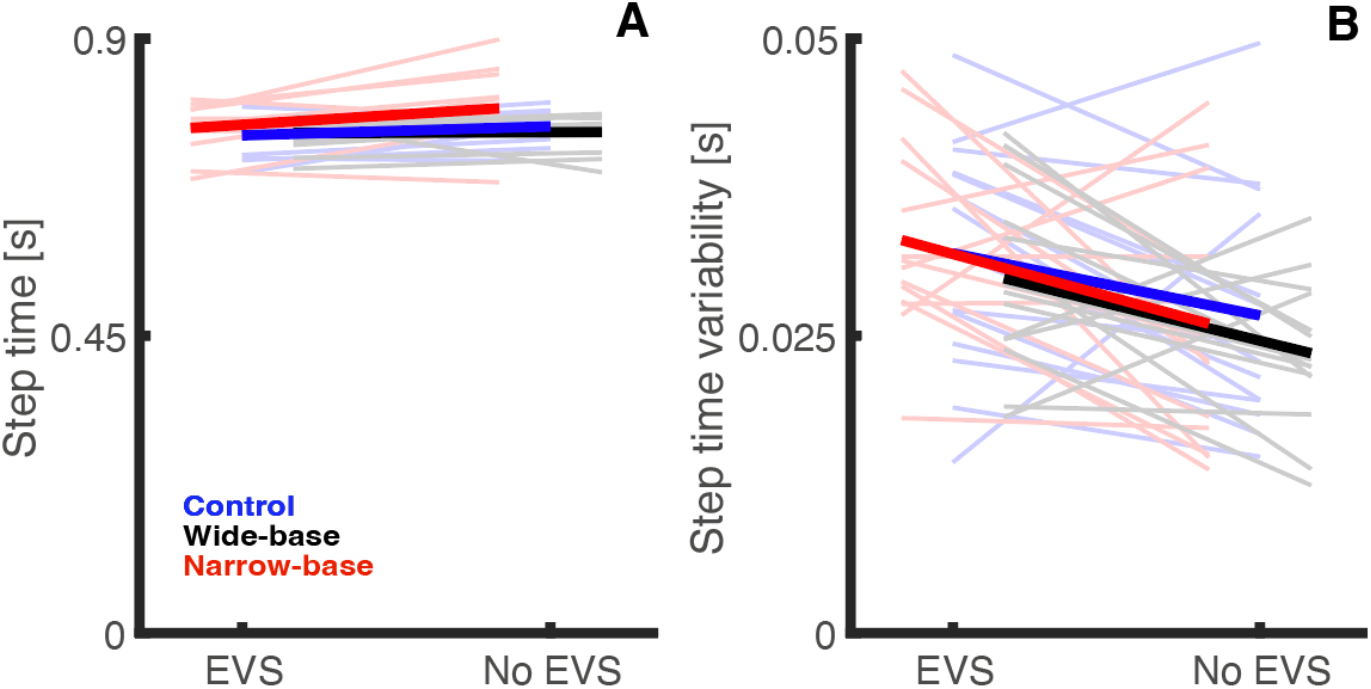
Effects of Condition and EVS on step time (A) and step time variability (B). Thick lines represent mean values and thin lines show individual data for each condition according to the following colors - blue lines: control condition; black lines: wide-base condition; red lines: narrow-base condition. Note: the conditions are jittered on the x-axis.

As hypothesized, the LDE and variability of CoM movement were (Figure 4) was significantly increased (i.e. decreased local dynamic stability) during EVS conditions. In addition, there was a significant effect of Condition on LDE. The post-hoc tests (Table 4) indicated that the narrow-base condition was more stable than both other conditions, while CoM variability was greater in narrow-base than in normal walking (Table 2). In contrast with our hypothesis, there was no significant interaction effect on LDE or variability.

**Figure 4:**
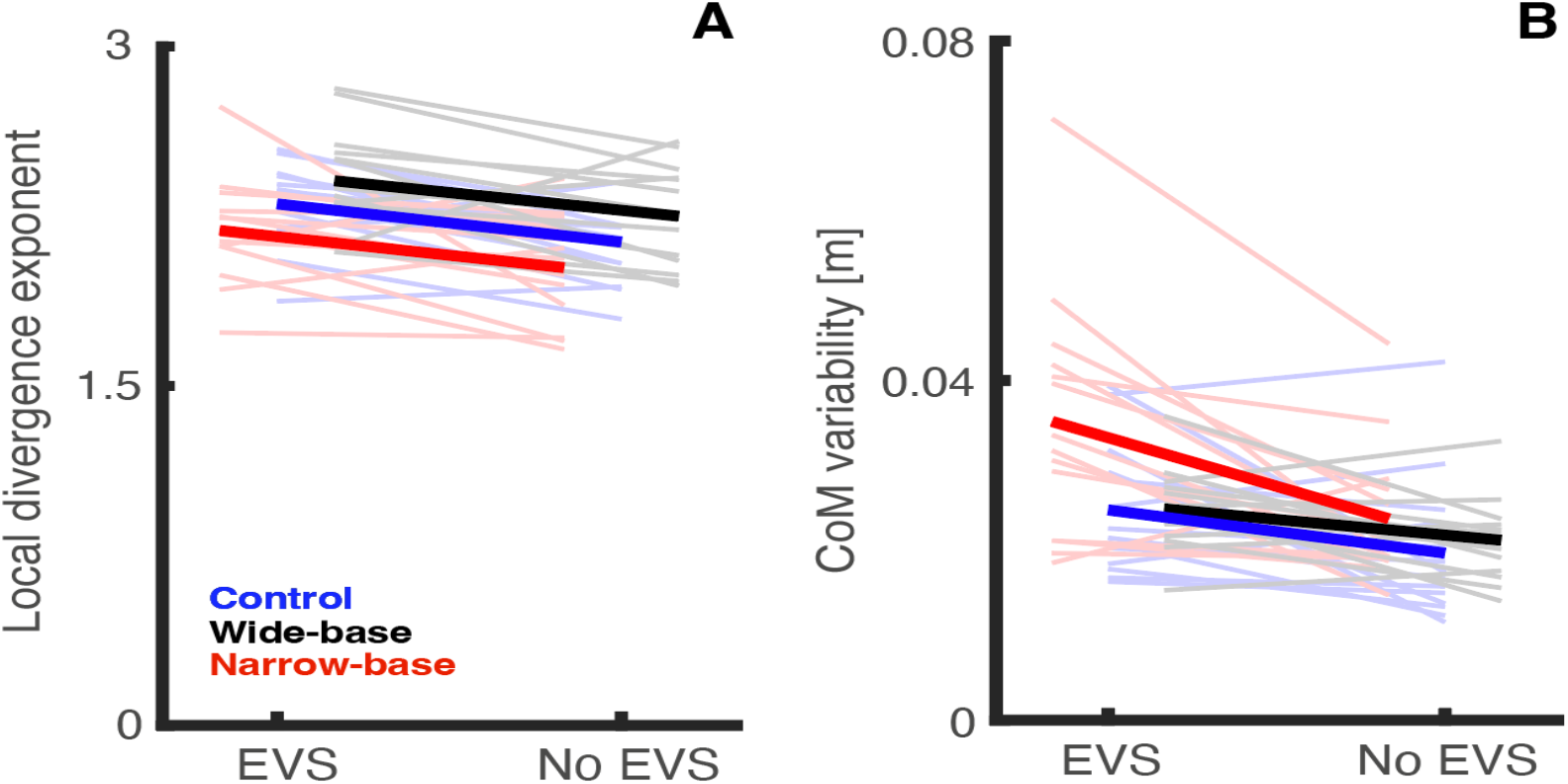
Effects of Condition and EVS on (A) mediolateral local divergence exponent and (B) variability of CoM position at heel strike (calculated from T6 marker velocity). Thick lines represent mean values and thin lines show individual data for each condition according to the following colors - blue lines: control condition; black lines: wide-base condition; red lines: narrow-base condition. Note: the conditions are jittered on the x-axis.

In contrast with our hypothesis, no main effect of EVS on R^2^ between CoM state and foot placement was found (Figure 5A). However, as hypothesized, residual variance was significantly higher with than without EVS (Figures 5B). Condition had a significant effect on R^2^ and residual variance. Post-hoc tests indicated that R^2^ and residual variance were greater in the narrow-base condition than in the control condition and only the R2 was higher in the narrow-base than the wide-base condition (Table 2).

**Figure 5:**
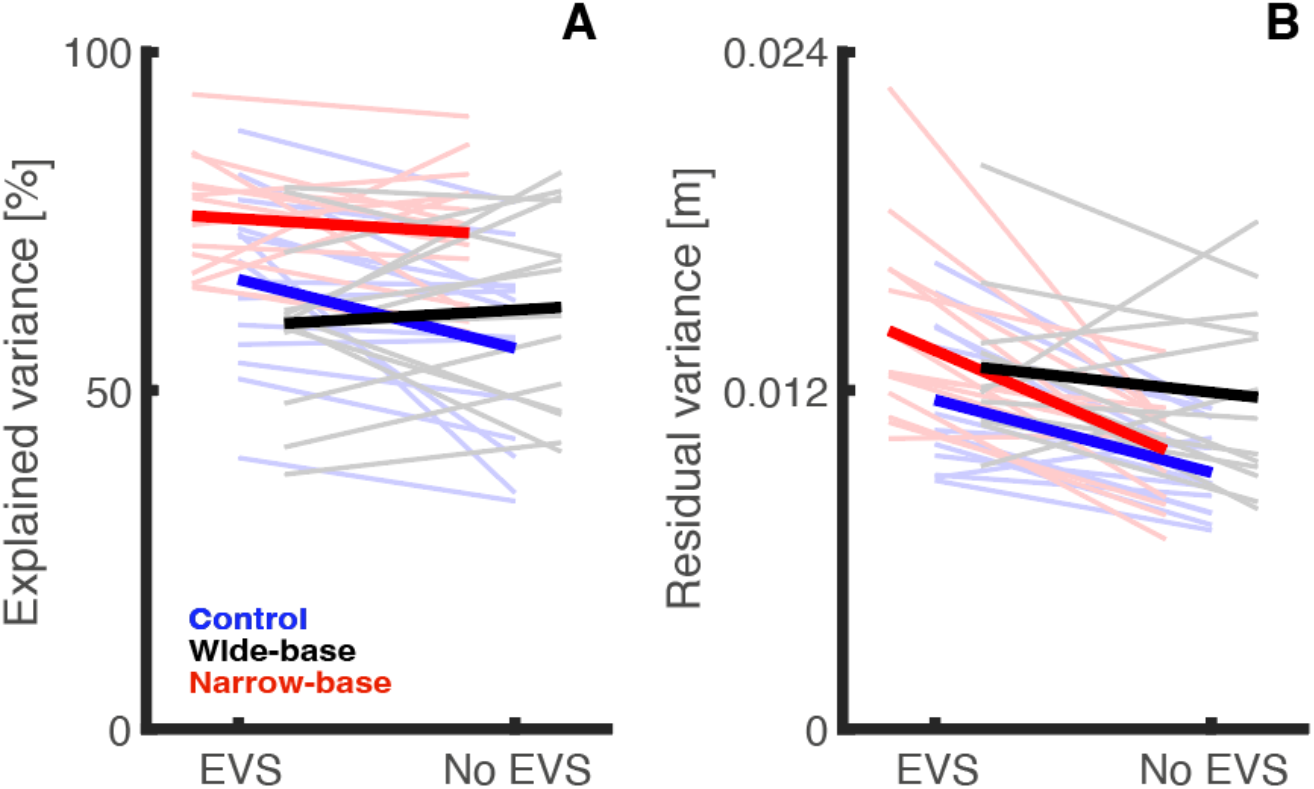
Effects of Condition and EVS on R^2^ (percentage of explained variance in foot placement) (A), the residual variance in foot placement (B) and center of mass position variability (C). Thick lines represent mean values and thin lines show individual data for each condition according to the following colors - blue lines: control condition; black lines: wide-base condition; red lines: narrow-base condition. Note: the conditions are jittered on the x-axis.

Significant interaction effects of EVS and Condition on R^2^ and residual variance were found. Post-hoc differences were found only between control and wide-base walking for R^2^ and between narrow-base and wide-base walking for the residual variance (Table 3). The former reflects a larger effect of EVS on R^2^ in the control condition compared to the wide-base condition, with an unexpected sign in the control condition. The latter findings is in line with our expectation of decreasing effects of EVS going from narrow-based to normal to wide-based walking.

## Discussion

We tested the effects of stochastic EVS on gait stability when walking with different step widths. As we had previously shown a decreased coupling between EVS and ground reaction forces in wide-base walking compared to control walking, we expected that EVS during wide-base walking would have less effect on gait stability than during control walking. Likewise, as we have shown increased coupling between EVS and ground reaction forces in narrow-base walking, we hypothesized a larger effect of EVS on gait stability during narrow-base walking than during control walking. While EVS was found to decrease gait stability and increase gait variability and the residual error in foot placement, the hypothesized interactions with walking condition were not consistently found.

### Effects of EVS

We hypothesized that stochastic EVS would decrease gait stability by perturbing feedback control of foot placement. Our results support this hypothesis. Walking with EVS led to reduced gait stability during walking at all step widths, evidenced by a higher LDE, as found previously (Sloot et al. 2011; van Schooten et al. 2011; Magnani et al. 2021). We found no significant effect of EVS on R^2^ a, and the figures even suggested an increase in R^2^ when walking with EVS for the control and narrow base conditions. This would falsify our hypothesis. However, as in previous studies, we also found that when walking with EVS, the variability of trunk movement increased (van Schooten et al. 2011), which could potentially explain the higher R^2^ values. Indeed, the residual variance in foot placement increased when walking with EVS, thus the remaining errors in foot placement were still larger, despite foot placement being more coupled to trunk motion.

Step width increased with EVS possibly as an adaptation to preserve stability while exposed to EVS (Hak et al. 2012). Despite walking to the beat of a metronome in all conditions, step time showed a slight decrease when walking with EVS. This indicates that stride frequency increased. Previous studies have also found small and significant increases in stride frequency when participants walked in challenging conditions in spite of the use of a metronome (van Leeuwen et al. 2020; Hoogstad et al. 2022). Similar to the increase in step width, an increase in stride frequency may serve as an adaptation to preserve gait stability (Hak et al. 2012).

### Effects of walking condition

Gait stability was largest in narrow-base walking and lowest in wide-base walking. This may reflect the need to more tightly control gait when the base of support is smaller. Feedback gains in foot placement control increase with decreasing average step width (Perry and Srinivasan 2017). Similarly, we found a higher R^2^ in narrow-base walking than in normal and wide-base walking. This may reflect that a decreased base of support decreases margins of safety, so it would need more precise foot placement modulation (Perry and Srinivasan 2017; van Leeuwen et al. 2020). In apparent contrast, we found that residual variance in foot placement was smallest during normal walking. Possibly the fact that this condition did not impose a constraint on foot placement, whereas both other conditions did accounts in part for this.

The larger residual error in foot placement in the narrow-base condition seems at odds with the fact that local dynamic stability was highest in this condition. However, it should be kept in mind that foot placement is only one mechanism by which stability can be controlled. Other mechanisms are shifting the center of pressure under the stance foot (Reimann et al. 2018; van Leeuwen et al. 2021) and changing angular momentum around the CoM (Hof 2008; van den Bogaart et al. 2020). The increase in EVS-erector spinae muscle coupling during narrow-base walking that we found previously (Magnani et al. 2021), suggests that indeed angular momentum changes may play a larger role during narrow-base walking, as was also seen in behavioral studies (Bogaart et al. 2021).

### Interaction effects of walking condition and EVS

We hypothesized that EVS effects on gait stability and foot placement would be stronger in narrow-base walking than in control walking and less strong in wide-base walking. An interaction was not found for gait stability but it was found for the residual error in foot placement. The latter result was in line with our hypothesis, with larger effects for narrow-base walking, and smaller effects for wide-base walking, although, post-hoc differences were mostly not significant. Thus, although vestibular contributions to control of foot placement may modulate with step width, this did not translate to different effects of a vestibular error signal on gait stability at different step widths. Possibly due to concomitant adaptations in average step width and step time.

### Limitations

This study has some limitations. First of all, our results were obtained during treadmill walking. Mechanically there are no differences between overground and treadmill walking (van Ingen Schenau 1980). Nevertheless, step width has been shown to be wider and step width variability was reduced on a treadmill compared to overground (Rosenblatt and Grabiner 2010), suggesting potential differences in control of treadmill vs. overground walking. Another limitation is that we used T6 as a proxy of the CoM, and this is a rather high location. Nevertheless, the R^2^ between CoM and foot placement was similar as in previous work using a more accurate estimate of the CoM or the pelvis as a proxy (Mahaki et al. 2019; van Leeuwen et al. 2021).

## Conclusion

In conclusion, stochastic EVS decreases local dynamic stability of gait, but this effect is not different when walking at different step widths. Residual variance in foot placement was increased with EVS, indicating less accurate foot placement. Still, this effect was not significantly different between different step widths. Thus, a vestibular error signal decreases gait stability, but this decrease is not significantly modulated by step width. This is different from our earlier work which showed that vestibular contributions to control of gait are modulated by stabilizing demands.

## Declarations

### Availability of data and material (data transparency)

The data and code for this manuscript can be found at https://surfdrive.surf.nl/files/index.php/s/dUu3uRQyVHlN2g3 and will be published to Zenodo upon acceptance.

